# Berberine and obatoclax inhibit SARS-CoV-2 replication in primary human nasal epithelial cells in vitro

**DOI:** 10.1101/2020.12.23.424189

**Authors:** Finny S. Varghese, Esther van Woudenbergh, Gijs J. Overheul, Marc J. Eleveld, Lisa Kurver, Niels van Heerbeek, Arjan van Laarhoven, Pascal Miesen, Gerco den Hartog, Marien I. de Jonge, Ronald P. van Rij

## Abstract

Severe acute respiratory syndrome coronavirus 2 (SARS-CoV-2) emerged as a new human pathogen in late 2019 and has infected an estimated 10% of the global population in less than a year. There is a clear need for effective antiviral drugs to complement current preventive measures including vaccines. In this study, we demonstrate that berberine and obatoclax, two broad-spectrum antiviral compounds, are effective against multiple isolates of SARS-CoV-2. Berberine, a plant-derived alkaloid, inhibited SARS-CoV-2 at low micromolar concentrations and obatoclax, originally developed as an anti-apoptotic protein antagonist, was effective at sub-micromolar concentrations. Time-of-addition studies indicated that berberine acts on the late stage of the viral life cycle. In agreement, berberine mildly affected viral RNA synthesis, but strongly reduced infectious viral titers, leading to an increase in the particle-to-pfu ratio. In contrast, obatoclax acted at the early stage of the infection, in line with its activity to neutralize the acidic environment in endosomes. We assessed infection of primary human nasal epithelial cells cultured on an air-liquid interface and found that SARS-CoV-2 infection induced and repressed expression of a specific set of cytokines and chemokines. Moreover, both obatoclax and berberine inhibited SARS-CoV-2 replication in these primary target cells. We propose berberine and obatoclax as potential antiviral drugs against SARS-CoV-2 that could be considered for further efficacy testing.

## Introduction

Coronaviruses form a group of respiratory viruses in the order *Nidovirales* that possess a positive-sense RNA genome of approximately 30 kb (1). Two coronaviruses of zoonotic origin caused significant outbreaks in the near past, severe acute respiratory syndrome coronavirus (SARS-CoV) (2, 3) and Middle Eastern respiratory syndrome coronavirus (MERS-CoV) (4). In late 2019, a new coronavirus closely related to SARS-CoV emerged in Wuhan, China, and rapidly spread across the globe. As of today, more than 60 million SARS-CoV-2 cases have been confirmed worldwide, but recent estimates from the World Health Organization suggest that that the total number may be more than 20 times higher due to asymptomatic and undetected cases (5) (WHO Coronavirus Disease Dashboard, https://covid19.who.int/).

SARS-CoV-2 infects epithelial cells in the nasal and oral cavities via binding of the SARS-CoV-2 spike protein to the ACE2 receptor (6). The spike glycoprotein requires cleavage by the protease TMPRSS2 to convert it into an active form allowing fusion at the plasma membrane (7, 8). An additional furin cleavage site in the spike protein makes it also amenable to cleavage by intracellular proteases (9, 10). In permissive cells lacking TMPRSS2, receptor binding is most likely followed by dynamin and clathrin-mediated endocytosis of the virus particle into endosomal compartments (11). Acidification of these compartments leads to activation of cathepsin-B/L proteases which prime the spike protein, initiating membrane fusion and release of the encapsidated viral RNA into the cytoplasm (12, 13). Upon nucleocapsid disassembly, the viral positive-sense RNA genome is translated into two open reading frames that encode for several non-structural proteins [reviewed in (14–16)]. These proteins constitute the machinery that replicates the viral RNA and transcribes subgenomic RNAs that code for viral structural proteins and several accessory proteins (15). Newly formed viral genomic RNA is coated with nucleocapsid proteins, which interact with the structural proteins, resulting in budding into and transit through the ER-Golgi network and release of mature viral particles through exocytosis (15, 16). The virus then spreads to the lower respiratory tract and infects alveolar type-II pneumocytes in the lungs, where a severe infection could lead to acute respiratory distress syndrome (6, 17).

A year into the COVID-19 pandemic, there has been an unprecedented fast development of vaccines (18), as well as study of antiviral of adjunctive host-directed therapy. Apart from dexamethasone (19), however, none has been proved effective. As preventive measures including vaccines are unlikely to curb the epidemic alone, there is a clear need for improved antiviral therapy. This could be applied either preventively, or after symptom development.

In this study, we assessed the plant-based alkaloid berberine and obatoclax, an anticancer drug proven safe in clinical trials, as possible antiviral candidates against SARS-CoV-2. Both berberine (BBR) (20–22) and obatoclax (OLX) (23, 24) have broad-spectrum antiviral activity against a range of different viruses including herpes simplex virus, influenza A virus, chikungunya virus and Zika virus. We show that these compounds are effective against two different isolates of SARS-CoV-2 at low micromolar concentrations with promising selectivity indices. Antiviral activity was observed in Vero E6 cells as well as in physiologically relevant nasal epithelial cells cultured on an air-liquid interface. We propose these compounds for further assessment as antiviral agents against SARS-CoV-2.

## Results

### Berberine and obatoclax inhibit SARS-CoV-2 replication

BBR and OLX have shown antiviral activity against viruses from multiple families (20–24). We therefore tested these compounds against SARS-CoV-2 in Vero E6 cells. A virus growth curve was performed to determine the optimal time point for the assay. At a low multiplicity of infection (MOI) of 0.01, SARS-CoV-2 titers peaked at 24 hours post-infection (hpi) (Fig. S1A). Robust infection of Vero E6 cells was confirmed through staining infected cells for the presence of dsRNA replication intermediates and the SARS-CoV-2 spike protein. dsRNA staining was observed in the perinuclear area, likely corresponding to the ER-Golgi network containing SARS-CoV-2 replication complexes (Fig. S1B). Spike protein expression was also detected in the perinuclear area where they are synthesized, but also on the outer periphery of the cells, probably from viruses exiting the infected cells.

Next, a dose response assay was carried out under these conditions with a series of concentrations of BBR and OLX, using 0.1% DMSO as a negative control. Plaque assay titrations of the viral supernatants showed that both BBR and OLX are effective against SARS-CoV-2 with 50% effective concentration (EC_50_) values at low micromolar and nanomolar concentrations (EC_50 BBR_ = 9.1 μM and EC_50 OLX_ = 67 nM, respectively; Fig. 1A and B). Cytotoxicity of these compounds was tested under the same conditions, but in the absence of SARS-CoV-2 infection. BBR was toxic only at the highest concentrations tested (50% cellular cytotoxicity, CC_50_ > 150 μM), resulting in a selectivity index of >16 (Fig. 1C), whereas the CC_50_ value for OLX was 7.8 μM (Fig. 1D), corresponding to a high selectivity index of 116 due to its low EC_50_ value. Overall, these results indicate that BBR and OLX show potent antiviral activity against SARS-CoV-2 with good selectivity indices.

**Fig 1.**
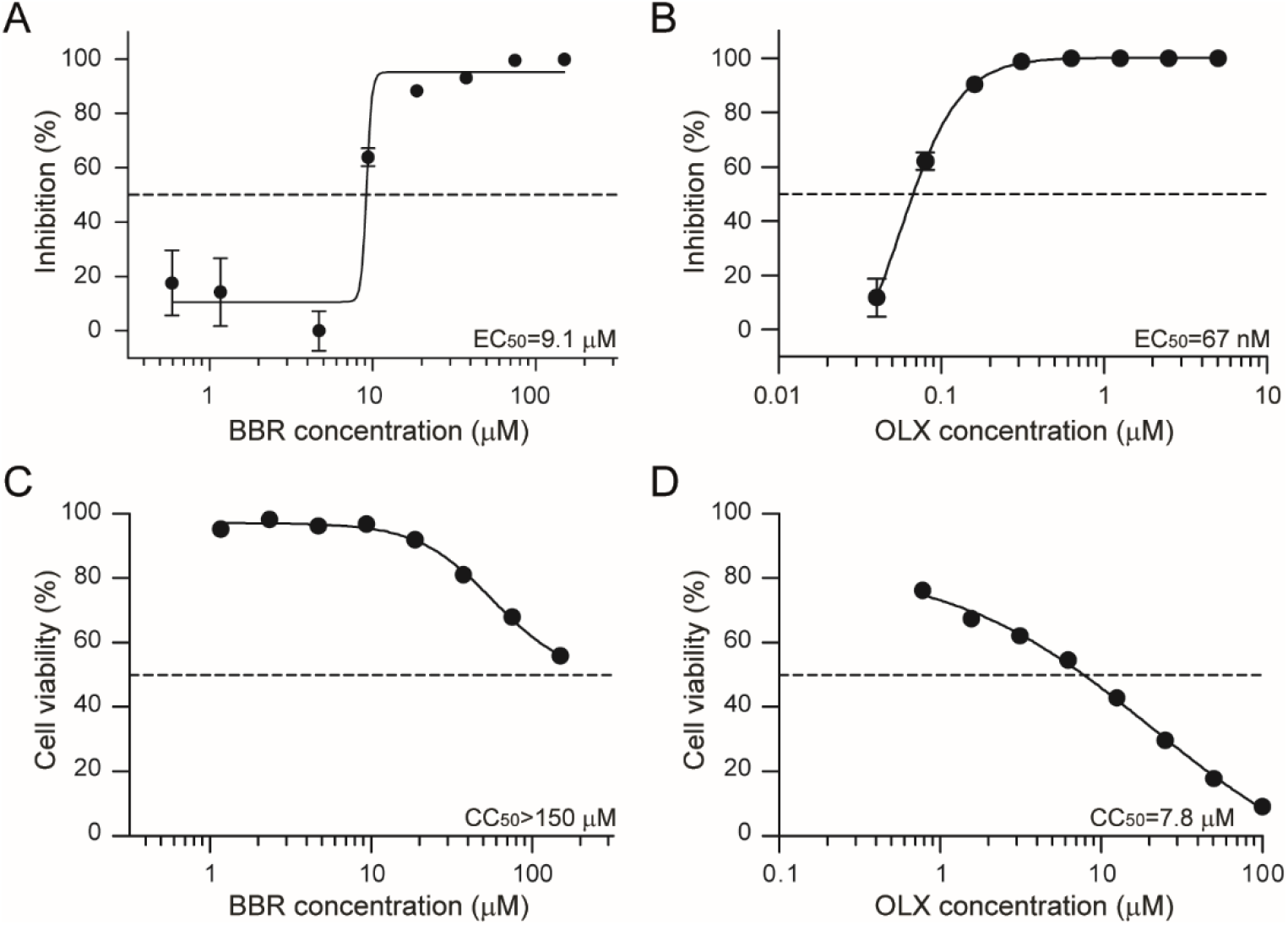
Berberine and obatoclax are effective antiviral compounds against SARS-CoV-2. Vero E6 cells were infected with SARS-CoV-2 (BavPat1 isolate) at an MOI of 0.01 for 24 h in the presence of the indicated concentrations of **(A)** berberine (BBR) or **(B)** obatoclax (OLX) in a two-fold dilution series or 0.1% DMSO. Infectious viral titers from duplicate cell culture supernatants were assessed by plaque assay and plotted as percentage inhibition compared to 0.1% DMSO control. Error bars indicate SD. Dashed line indicates 50% inhibition. A representative of two independent experiments is shown. Vero E6 cells were treated with the indicated concentrations of **(C)** berberine or **(D)** obatoclax or 0.1% DMSO control. 24h post-treatment, ATP content in cells from triplicate wells was measured as indication for cell viability, plotted as percentage compared to the DMSO control. Error bars indicate SD. A representative of two independent experiments is shown.

### Berberine and obatoclax are effective against a SARS-CoV-2 isolate from a different geographic region

Our initial antiviral assays were performed using a SARS-CoV-2 isolate from Bavaria, Germany (BavPat1). To assess whether BBR and OLX are also effective against another SARS-CoV-2 isolate, we isolated a SARS-CoV-2 isolate from a hospitalized COVID-19 patient at Radboud University Medical Center, the Netherlands. This isolate, SARS-CoV-2/human/Nijmegen/1/2020 (hereafter called Nijmegen1), had a relatively small plaque phenotype (Fig. S2), in agreement with the relatively passage history in Vero cells (25, 26). We used the Nijmegen1 isolate to perform dose response assays with BBR and OLX, using viral RNA in the cell culture supernatants as a readout (Fig. 2). OLX showed stronger antiviral activity for the Nijmegen1 isolate (EC_50_ = < 0.04 μM) (Fig. 2B) as compared to the BavPat1 isolate. In contrast, BBR showed relatively weak antiviral activity against the Nijmegen1 isolate (EC_50_ = 23.2 μM for viral RNA in the supernatant and EC_50_ = 43.3 μM for intracellular viral RNA; Fig. 2A) as compared to the initial experiments in which infectious titers were used as a readout. We therefore used conventional plaque assay to assess the antiviral activity of BBR against the SARS-CoV-2 Nijmegen1 isolate. We observed strong antiviral activity (EC_50_ = 2.1 μM) (Fig. 2C), which was slightly lower than for the BavPat1 isolate.

**Fig 2.**
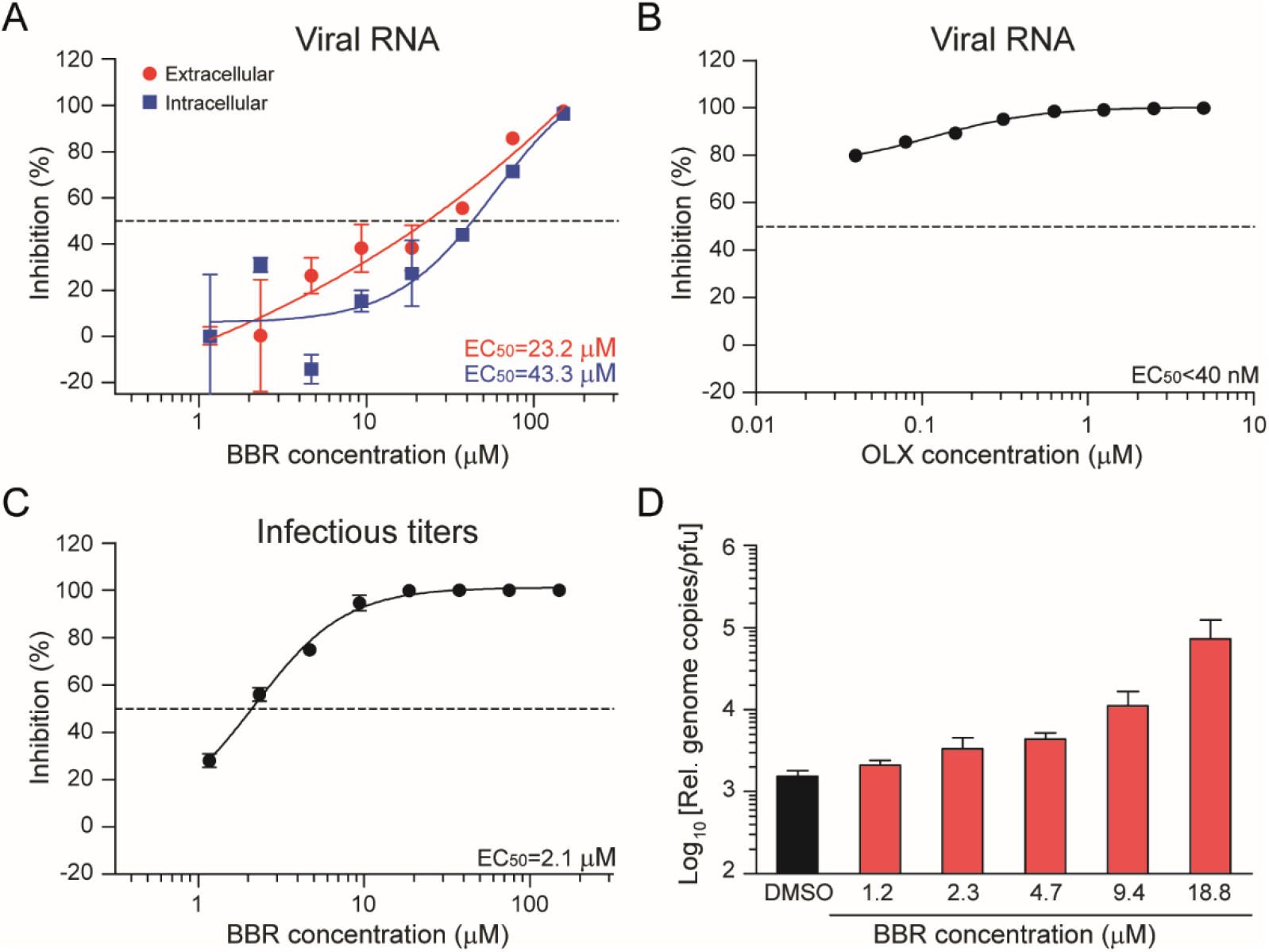
Berberine and obatoclax are effective against SARS-CoV-2 Nijmegen1 isolate. Vero E6 cells were infected with the SARS-CoV-2 Nijmegen isolate at an MOI of 0.01 for 24h in the presence of the indicated concentrations of **(A, C)** berberine (BBR) or (**B**) obatoclax (OLX) in a two-fold dilution series or 0.1% DMSO. **(A, B)** Viral RNA from duplicate cell culture supernatants (red line in panel A) was quantified by qRT-PCR and plotted as percentage inhibition compared to the DMSO control. For BBR-treated cells, both extracellular and Intracellular viral RNA was analyzed (red and blue lines, respectively). Intracellular viral RNA levels were normalized to the human β-actin housekeeping gene. **(C)** Infectious viral titers from duplicate cell culture supernatants were quantified by plaque assay and plotted as percentage inhibition compared to the DMSO control. Error bars indicate SD. Dashed line indicates 50% inhibition. **(D)** Ratio of relative genome copies to infectious viral particles in cell culture supernatants for the indicated, non-toxic berberine concentrations and DMSO control.

These results suggest that BBR does not inhibit viral RNA replication per se, but that it affects the production of infectious virus. To quantify this, we calculated the relative genome copy to infectious virus particle ratio for the different concentrations of berberine. Indeed, when compared to the DMSO control, this ratio increased upon BBR treatment in a dose-dependent manner (Fig. 2D). Altogether these results confirmed that both BBR and OLX are effective against a second, early passage isolate of SARS-CoV-2. Moreover, we conclude that BBR does not affect viral RNA replication or secretion of viral particles, but that it strongly affects the infectivity of the virus particles produced.

### Berberine and obatoclax act at different stages of the SARS-CoV-2 life cycle

To decipher a putative mode of action for BBR and OLX, we performed a time-of-addition assay. Vero E6 cells were infected at MOI of 1, adding 20 μM BBR or 0.25 μM OLX at different time points during the course of the experiment (Fig. 3A). Cell culture supernatants were harvested at 10 hpi and infectious viral titers were measured by plaque assay. OLX showed a very potent inhibition of 3 logs at the early stages of the viral life cycle. The levels of inhibition gradually decrease as OLX is added at later time points of the experiment (Fig. 3B), suggesting that OLX most likely affects the early stages of the SARS-CoV-2 infection cycle. Indeed, when OLX was added during the 1 h of virus adsorption and subsequently discontinued, it reduced viral titers by 1.5 logs, indicating that the compound inhibits viral entry.

**Fig 3.**
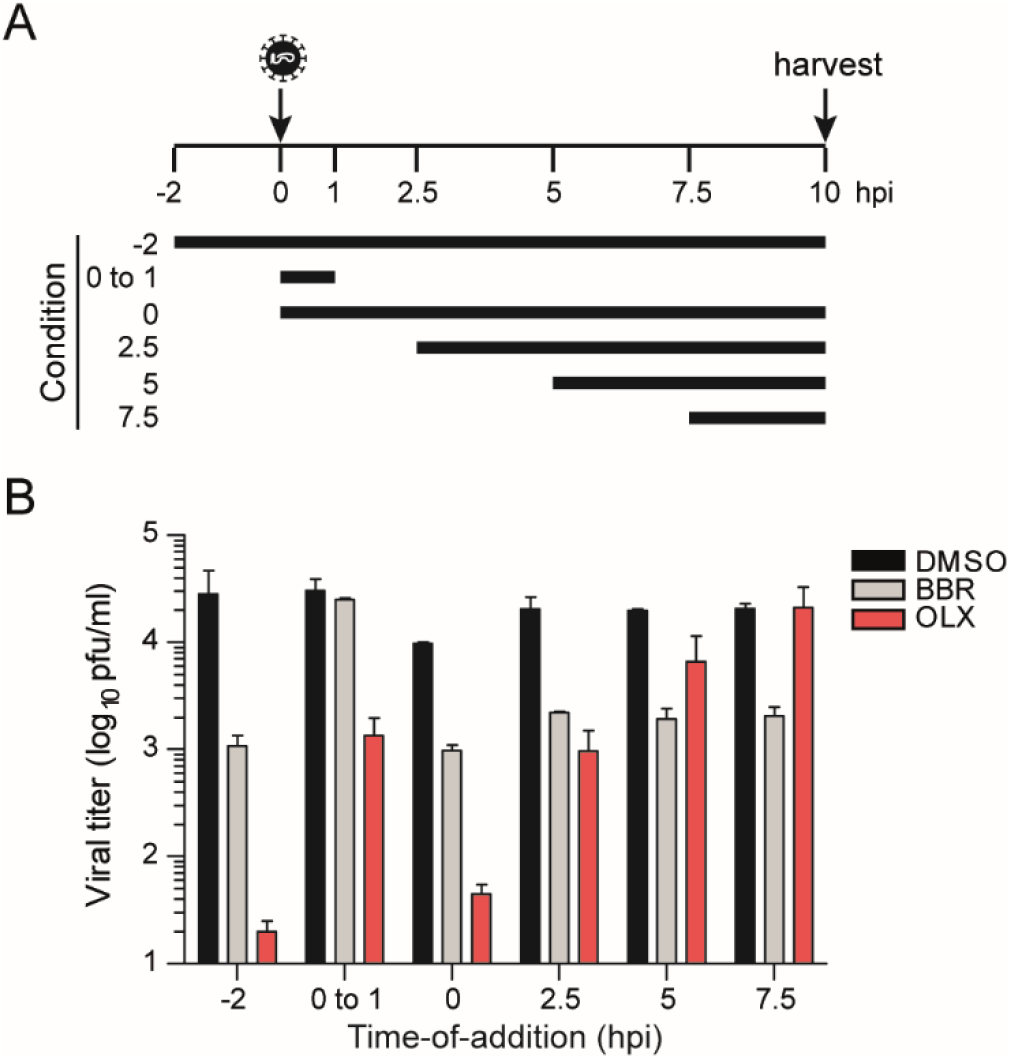
Time-of-addition assay. Vero E6 cells were infected with SARS-CoV-2 (BavPat1 isolate) at an MOI of 1. **(A)** Schematic layout of the assay. 20 μM berberine or 0.25 μM obatoclax or 0.1% DMSO was added to the infected cells at the indicated time points. **(B)** Plaque assay titers from cell culture supernatants collected at 10 hpi. Bars and error bars represent means and SD of n = 2 replicates.

For BBR, an inhibition of almost 1.5 logs was seen when the compound was added to the cells 2 h prior to inoculation and maintained throughout the experiment. However, no inhibition was observed when BBR is only added during the 1 h of virus adsorption. Potent inhibition of 1 log was observed when BBR was added during inoculation and continuously maintained thereafter. Strikingly, BBR continues to be equally effective at reducing viral titers even when added at 7.5 hpi, indicating that it acts late in the viral infection cycle (Fig. 3B), in line with our conclusion that BBR affects the production of infectious virus particles (Fig. 2).

### SARS-CoV-2 infection of primary nasal epithelial cells

Even though SARS-CoV-2 efficiently replicates in Vero E6 cells, these cells are derived from the African green monkey kidney and have a defective interferon pathway (27) and are thus not representative of natural target cells. It is therefore essential to validate candidate antiviral compounds in human host cells (28). We therefore cultured and differentiated human nasal epithelial cells, which were residual after surgery, on an air-liquid interface. This approach to mimic the natural environment of the infected host has previously been used in the study of circulating seasonal human coronaviruses, such as HCoV-229E, HCoV-OC43, HCoV-NL63, and HCoV-HKU1, as well as the zoonotic SARS and MERS coronaviruses (29).

Virus replication was assessed to determine the optimal time point for antiviral assays. We analyzed intracellular RNA, extracellular RNA from apical and basal supernatants, infectious viral titers and viral protein expression at different time points post-infection. Peak intracellular viral RNA levels were detected at 72 hpi (Fig. 4A). No time-dependent increase in viral RNA levels was observed in basolateral supernatants, suggesting inefficient release of virus into this compartment. In contrast, viral RNA signals from the apical surface supernatants were at their highest at 72 hpi, corresponding to the kinetics of intracellular RNA (Fig. 4B). Preferential release from the apical membrane in polarized epithelial cells has been observed before for multiple viruses, including SARS-CoV (30–32). Infectious viral titers from the apical supernatants also showed active viral replication, with peak titers at 72 hpi (Fig. 4C) in line with the viral RNA kinetics (Fig. 4A, 4B).

**Fig 4.**
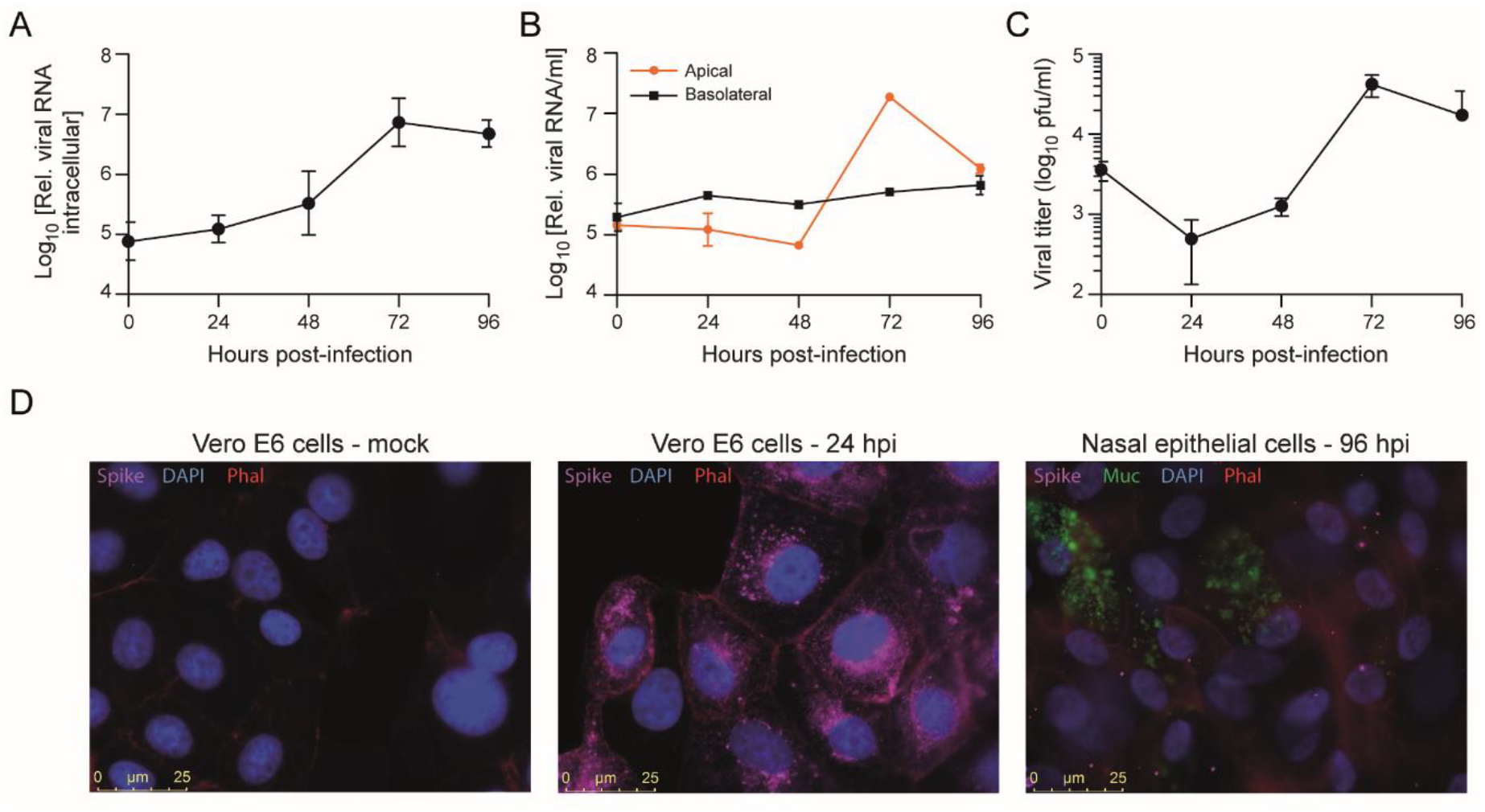
SARS-CoV-2 infection of primary nasal epithelial cells. Primary nasal epithelial cells, cultured on an air-liquid interface, were infected with SARS-CoV-2 (BavPat1 isolate) at an MOI of 10. At the indicated time points, cells were harvested and viral RNA from **(A)** cells or **(B)** apical and basolateral compartment supernatants was quantified by RT-qPCR. Intracellular viral RNA levels were normalized to the human β-actin housekeeping gene. Error bars indicate SD (n=2). **(C)** Infectious viral titers from apical supernatants corresponding to the indicated time points were quantified by plaque assay. Error bars indicate SD (n=2). **(D)** Immunofluorescence staining of mock (left panel), SARS-CoV-2 infected Vero E6 cells fixed at 24 hpi (MOI 0.01; middle panel) or SARS-CoV-2 infected primary nasal epithelial cells fixed at 96 hpi (MOI 10; right panel). The presence of SARS-CoV-2 Spike protein subunit S1 (pink) was assessed in cells stained with DAPI (nucleus, blue) and phalloidin (F-actin, red). Nasal epithelial cells were additionally stained with anti-Muc5AC antibodies (goblet cells, green). Bar represents 25 μm.

The nasal epithelium contains both ciliated cells and mucus producing goblet cells after 4 weeks of differentiation on the air-liquid interface (Fig. S3). To assess infection of these cell types, we analyzed SARS-CoV-2 spike protein expression by immunofluorescence microscopy. To validate the anti-Spike protein antibody used, we first analyzed SARS-CoV-2 infected Vero E6 cells and observed abundant S protein expression in the perinuclear area as well as the periphery (Fig. 4D). In SARS-CoV-2 infected nasal epithelial cells, SARS-CoV-2 S protein was detected from 48-72 hpi onwards (Fig. S3). In these cells, the S protein was mostly located at the outer cell periphery, and, as expected, at much lower levels than in Vero E6 cells. At 0 hpi, some S protein staining is also visible, likely corresponding to viral particles that have not internalized and remain attached to the cell surface (Fig. S3). Additionally, the epithelial cells were stained for the differentiation markers tubulin IV and Muc5Ac to identify ciliated cells and goblet cells, respectively. It did not appear that viral infection was more abundant in either of these cell types (Fig. 4D, S3).

### Cytokine and chemokine responses are selectively enhanced or inhibited during SARS-CoV-2 infection of primary nasal epithelial cells

To assess the epithelial (immune) response to SARS-CoV-2 infection, we measured cytokine and chemokine concentrations in the cell culture medium of primary nasal epithelial cells infected with SARS-CoV-2 from 0 to 96 hpi. Cytokines and chemokines associated with granulocyte (CXCL1 and CXCL8), monocyte (CXCL10, CCL4, and CCL20), NK cell (CXCL10 and CCL4) and T cell (CCL20, CCL17, and CXCL10) chemotaxis and activation increased over time compared to the mock infected cells (Fig. 5A). CXCL1, CXCL8, CXCL10, CCL4, and CCL20 showed a more rapid production rate between 48 and 96 hpi, which is in line with the viral replication kinetics that we observed (Fig. 4), suggesting that viral replication triggers cytokine and chemokine production in these cells. Strikingly, the cytokines that are typically associated with viral infections and acute phase reactions (IL-1β, TNF-α2, IL-6, IFN-λ1, and IFN-λ2,3) did not show increased production compared to mock infected cells, with the exception of IL-1β (Fig. 5A and Fig. S4). Furthermore, within the first 48 h of infection, the production of CXCL5, CCL3, CCL5, and, to a lesser extent, GM-CSF was reduced (Fig. 5B). This could be indicative of immune evasive activity of SARS-CoV-2.

**Fig. 5.**
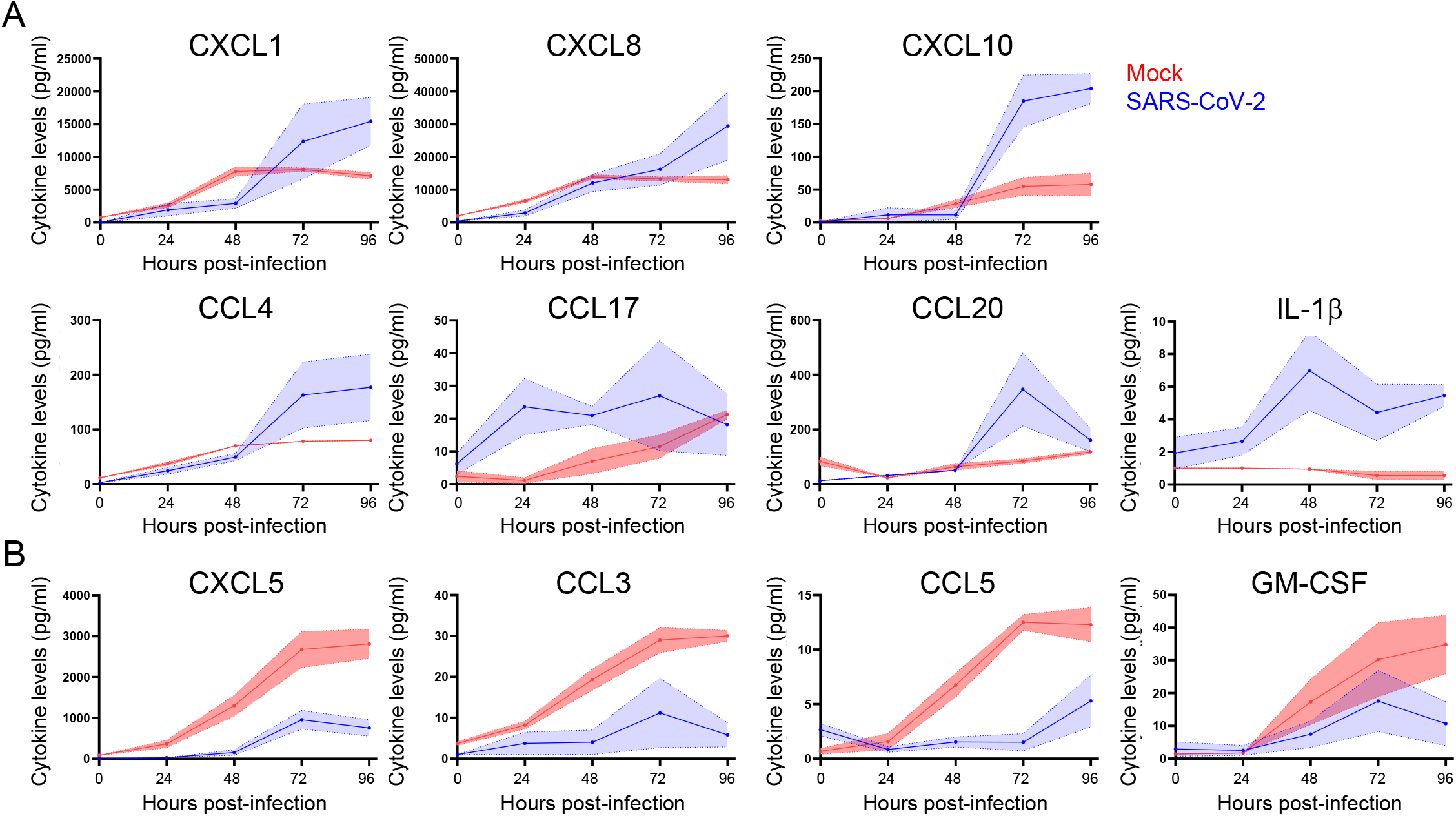
Induction and repression of specific cytokine and chemokines during SARS-CoV-2 infection of primary nasal epithelial cells. Primary nasal epithelial cells cultured on an air-liquid interface were infected with SARS-CoV-2 (BavPat1 isolate) at an MOI of 10 or mock infected. At the indicated time points, medium from the basolateral compartment was harvested and the concentration of the indicated cytokines and chemokines was analyzed by a bead-based immunoassay. Cytokines and chemokines with increased (**A**) and decreased (**B**) expression are shown. Means and SEM (shading) of n = 2 replicates are shown.

These data suggest that the epithelium selectively stimulates cellular immune responses in response to SARS-CoV-2 infection, while the virus inhibits secretion of specific cytokines. Both processes appear to be selective as for various other cytokines no difference in production could be observed between infected cells and mock infected cells (Fig. S4). Other cytokines that we tested, such as IFN-α, IFN-β, IL-10, and IL-12p70 were not produced by the epithelium at all (data not shown). Together, these data indicate a specific and dynamic interaction between airway epithelial cells and SARS-CoV-2.

### Berberine and obatoclax are effective against SARS-CoV-2 in nasal epithelial cells

Having established primary nasal epithelial cells as a relevant cellular model for SARS-CoV-2 infection, we assessed the antiviral activity of BBR and OBX in these cells. Nasal epithelial cells were infected with SARS-CoV-2 in the presence of increasing concentrations of BBR or OLX along with DMSO as a negative control and viral RNA levels were assessed at 72 hpi. BBR was effective at inhibiting SARS-CoV-2 RNA levels in the supernatant of this nasal epithelial cell model with an EC_50_ value of 10.7 μM (Fig. 6A), similar to that seen in Vero E6 cells. Likewise, OLX was effective in the sub-micromolar range with an EC_50_ value of 0.2 μM. Cytotoxicity assays were done with BBR and OLX over a 72h time period. There was a slight increase in toxicity in these cells as compared to the shorter 24 h exposure period in Vero E6 cells (Fig. 6C and D), resulting in lower selectivity indices for both compounds (SI=8.1 and 32.7 for BBR and OLX, respectively) than in Vero E6 cells. Altogether, these results indicate that BBR and OLX effectively inhibit SARS-CoV-2 replication in a physiologically relevant human nasal epithelial cell model.

**Fig 6.**
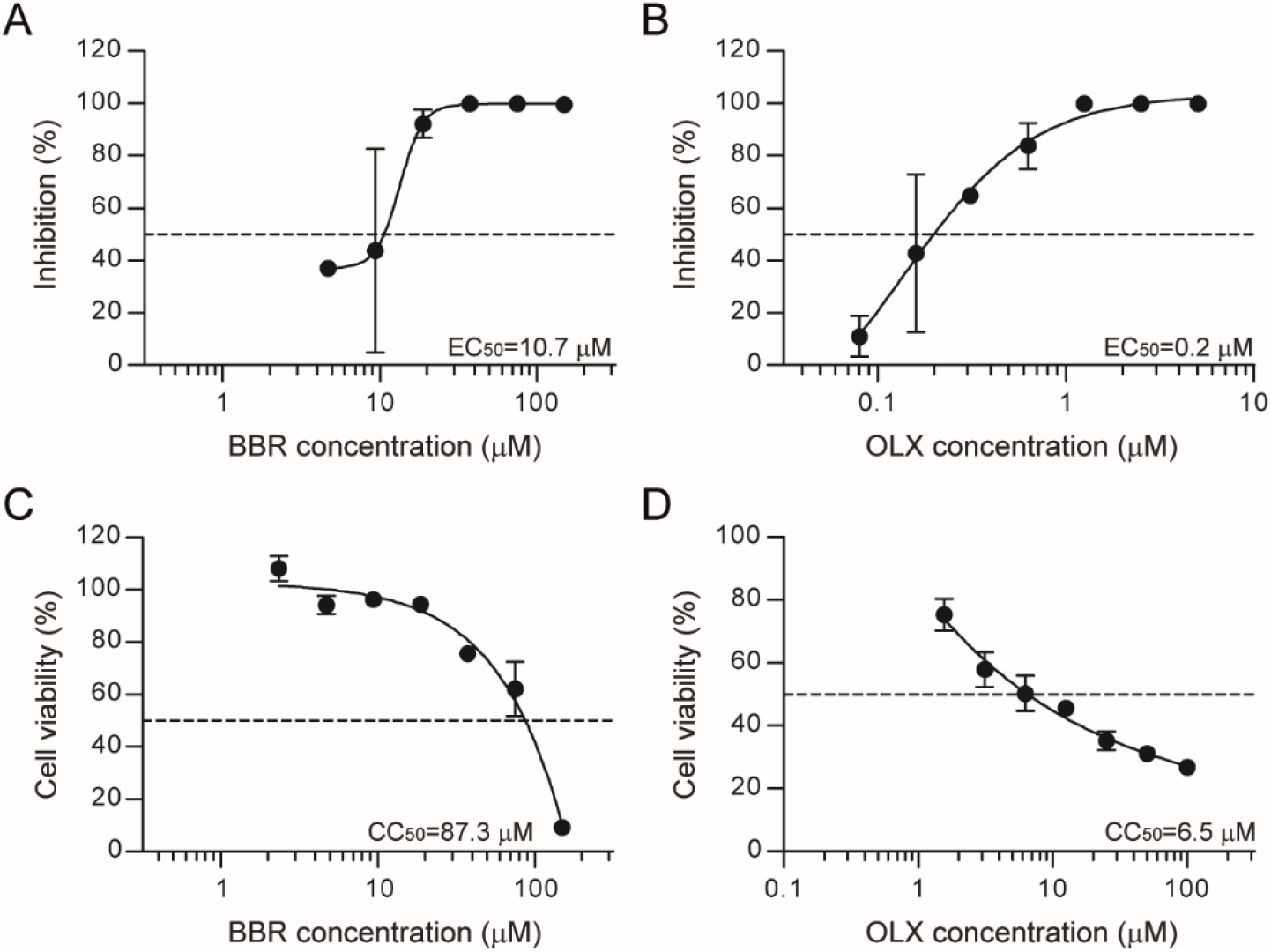
Berberine and obatoclax inhibit SARS-CoV-2 in primary nasal epithelial cells. Primary nasal epithelial cells cultured on an air-liquid interface were infected with SARS-CoV-2 (BavPat1 isolate) at an MOI of 10 in the presence of the indicated concentrations of **(A)** berberine (BBR) or **(B)** obatoclax (OLX) in a two-fold dilution series or 0.1% DMSO. Viral RNA from duplicate apical surface supernatants at 72 hpi was quantified by qRT-PCR and plotted as percentage inhibition compared to the DMSO control. Error bars indicate SD (n=2). Dashed line indicates 50% inhibition. Primary nasal epithelial cells cultured on an air-liquid interface were treated with the indicated concentrations of **(C)** berberine or **(D)** obatoclax or 0.1% DMSO control. At 72 h, ATP content in cells from duplicate samples was measured as a measure of cell viability, plotted as percentage of the DMSO control. Error bars indicate SD. Dashed line indicates 50% survival. A representative of two independent experiments is shown.

## Discussion

The COVID-19 pandemic has rapidly spread across the world, causing large-scale morbidity and mortality (5) (WHO Coronavirus Disease Dashboard, https://covid19.who.int/) and huge economic losses (33). Effective treatment for patients who fall ill despite vaccination or other preventive measures is urgently needed. In this study, we propose to repurpose BBR and OLX, two compounds with good safety profiles in clinical trials as potential therapeutic options against SARS-CoV-2.

BBR is an isoquinoline alkaloid derived from the Chinese herb *Coptis chinensis* and plants of the *Berberis* genus (34). Its wide-ranging biological properties identified in pre-clinical studies include anti-inflammatory, anti-arrhythmic, antimicrobial (35), and cholesterol-lowering (36) activity. In a large phase IV study randomising 612 patients, a treatment regimen containing 1000 mg BBR daily was not more effective in treating *Helicobacter pylori* than a comparator treatment, but it was well tolerated apart from a bitter taste (20%) and nausea (12%) as the most outspoken side effects (37). Previous studies also identified a favourable safety profile, with predominantly mild gastrointestinal side effects (38). BBR has not found a place in clinical practise, but five placebo-controlled clinical trials are currently recruiting patients to study the effect of BBR against a range of conditions, including colorectal adenomas, schizophrenia and diabetes (ClinicalTrials.gov Identifiers NCT03281096, NCT03333265, NCT03378934, NCT02808351, NCT02983188, NCT03976336, NCT03198572, NCT02737943).

BBR has broad spectrum antiviral activity *in vitro* against viruses from several different families, including influenza A virus, enterovirus, chikungunya virus, hepatitis B and C viruses, HIV, respiratory syncytial virus, human cytomegalovirus, herpes simplex virus and human papilloma virus [reviewed in (39)]. We show here that BBR is effective against SARS-CoV-2 at low micromolar concentrations in Vero E6 cells. While this manuscript was in preparation, our results were confirmed by another study showing the antiviral activity of BBR against SARS-CoV-2 also in Vero E6 cells at a similar EC_50_ value (10.6 μM) (40). Our study adds the validation of this finding in physiologically relevant primary nasal epithelial cells, where BBR is equally effective against SARS-CoV-2. Furthermore, we show that BBR acts late in the viral life cycle and likely induces the formation of non-infectious virus particles. Congruently, BBR also inhibited replication of the alphaviruses Semliki Forest virus and chikungunya virus at a late stage (41), likely by interfering with capsid protein-viral RNA interactions, leading to nucleocapsid assembly or disassembly defects and less infectious viral particles (42). It remains to be confirmed whether BBR has similar effects on assembly of SARS-CoV-2 virions. In addition to inducing possible assembly affects, BBR may affect cellular pathways to reduce SARS-CoV-2 replication. BBR targets several cellular signalling pathways including major MAP-kinase pathways (ERK, p38 MAPK and JNK), as well as the NF-κB and the AMPK/m-TOR signalling pathways (43, 44). Many viruses upregulate these pathways to maximally utilize cellular resources for their replication (45). The modulation of these pathways by BBR has been implicated in its antiviral activity against several different viruses, including chikungunya virus (22), enterovirus 71 (21) and herpes simplex virus (20). A recent proteomic analysis of SARS-CoV-2 interaction networks revealed that the p38 MAP kinase pathway was upregulated to counteract the host immune system. In agreement, inhibitors specifically targeting the p38 MAPK displayed antiviral activity against SARS-CoV-2 (46). Modulation of these cellular pathways needed for productive virus infection, could be a second mechanism through which BBR inhibits SARS-CoV-2 replication.

OLX was originally developed as an inhibitor of Mcl-1, a member of the Bcl-2 family of anti-apoptotic proteins (47). OLX has been through phase I and II trials for a range of malignancies. Convincing antitumor effects have thus far not been demonstrated, but obatoclax proved safe in doses of 20-40 mg/m^2^. The most reported adverse events are dose-related grade 1 or 2 neurological symptoms shortly after infusion (48–53)

OLX has broad-spectrum antiviral activity, employing a mechanism independent of its pro-apoptotic activity. OLX acts as a weak base and rapidly neutralizes the acidic environment of endosomes and endolysosomes (23). The acidification of endosomes is employed as an entry mechanism by enveloped viruses of several different families, including SARS-CoV-2 in TMPRSS2-negative target cells where endosomal cathepsin B/L proteases are activated by a drop in endosomal pH (7, 13, 54). We found that OLX inhibited SARS-CoV-2 at sub-micromolar concentrations in Vero E6 cells (EC_50_ = 0.06 μM) with a very high selectivity index. This confirms the results of another study in which OLX was reported to inhibit SARS-CoV-2-induced cytotoxicity in Vero E6 cells (55). Our time-of-addition assays confirmed the early effect of OLX, in alignment with the expected mechanism of virus entry inhibition. Moreover, we observe antiviral activity in nasal epithelial cells, with a three-fold higher EC_50_ value as compared to Vero E6 cells. This is probably due to SARS-CoV-2 entry in airway epithelial cells being predominantly dependent on TMPRSS2 protease priming of the viral spike protein (7, 54). However, unlike chloroquine, which also increases the endosomal pH but does not inhibit SARS-CoV-2 replication in lung epithelial cells (28), we observe an antiviral effect with OLX albeit with a lower but acceptable selectivity index. In clinical studies where OLX was administered intravenously over 3 hours, plasma level concentrations of ~0.4 μM were obtained (48, 49), suggesting that it would be possible to achieve therapeutic dose levels to treat COVID-19.

To identify candidate antiviral compounds, it is important to assess antiviral activity in physiologically relevant *in vitro* models as drug sensitivities may be cell-type specific (28, 56). We therefore characterized SARS-CoV-2 infection in primary nasal epithelial cells, which are one of the first cells the virus encounters during infection (57). Expectedly, in these cells with an active innate immune system, SARS-CoV-2 showed lower replication kinetics and reached lower titers than in Vero E6 cells. Still, robust time-dependent increase in SARS-CoV-2 replication was seen using multiple assessment parameters – viral RNA levels, titers and viral protein expression. The viral titers we obtained correspond well to those observed previously in human tracheobronchial epithelial cells grown on an air-liquid interface (58). In undifferentiated human airway epithelium, reconstituted from human primary nasal cells from a pool of 14 donors, higher SARS-CoV-2 titers were observed (59). We used differentiated cells from the post-operative tissue of a single donor, which may contribute to the observed differences.

SARS-CoV-2 infection of primary nasal epithelial cells induced epithelial release of CXCL1, CXCL8, CXCL10, CCL4, CCL20, CCL17 and IL1β. In contrast, we did not observe an increase in TNFα, IL-6, IFN-β, IFN-λ1, and IFN-λ2,3, cytokines that are typically involved in antiviral reactions and acute phase induction (60, 61). These observations might suggest that epithelial cells recruit leucocytes to the site of infection, rather than initiating an acute phase reaction themselves. Interestingly, viral infection also leads to reduced production of CXCL5, CCL5, CCL3 and GM-CSF, which are mainly involved in the attraction and activation of granulocytes and monocytes (62), suggesting that SARS-CoV-2 actively inhibits the recruitment of leukocytes early in infection.

In contrast to our findings, Pizzorno *et al.* observed a minor increase in IL-1β and TNFα mRNA levels, and a larger increase in IFN-λ1 and IFN-λ2,3,4 mRNA levels in nasal epithelial cells infected with SARS-CoV-2 (59). This discrepancy could be due to several differences in experimental setup, including the readouts (intracellular RNA versus protein in the supernatant), the use of cells from a pool of donors from different anatomical locations, and differentiation state (59).

In conclusion, our study puts forth two small molecule compounds, BBR and OLX, as repurposed antiviral molecules against SARS-CoV-2, in doses lower than previously shown to be safe in human clinical trials. While OLX is effective at early steps of the viral life cycle, likely interfering with entry processes, BBR acts on the later stages and likely reduces the infectivity of newly produced virions. BBR and OLX are effective in a physiologically relevant cell culture model at low micromolar concentrations and could be considered for further assessment.

## Materials and Methods

### Cells

African green monkey Vero E6 (ATCC CRL-1586) and Vero FM (ATCC CCL-81) kidney cells were grown in Dulbecco’s modified Eagle medium containing 4.5 g/L glucose and L-glutamine (Gibco), supplemented with 10% fetal calf serum (FCS, Sigma Aldrich), 100 μg/ml streptomycin and 100 U/ml penicillin (Gibco). Cells were maintained at 37°C with 5% CO_2_.

Post-operative residual tissue originating from the posterior nasal septum of a 17-year-old male was obtained after informed consent, according to the principles of the declaration of Helsinki. The patient had a fully normal nasal epithelium and no history of allergy, (chronic) rhinosinusitis or other mucosal disease. A single cell suspension was made by incubating the tissue in Hank’s Balanced Salt Solution (Gibco) with 1 mg/ml collagenase from clostridium histolyticum (Sigma) and 0.02 mg/ml DNase 1 (Sigma-Aldrich) for 2 h at 37°C. The cell suspension was filtered through a 70 μm cell strainer and the flow-through was collected. Cells were cultured and expanded in PneumaCult-Ex Basal plus Medium with supplements (Stemcell), 0.48 μg/ml hydrocortisone (Stemcell), and 100 μg/ml streptomycin and 100 U/ml penicillin (Lonza), according to manufacturer’s protocol (Stemcell). 50,000 primary nasal epithelial cells were seeded on 0.4 μm pore polyester membrane inserts (Corning) and expanded for 1 week in PneumaCult-Ex Basal plus Medium at 37°C and 5% CO_2_. When the cells had formed a tight epithelial layer, they were cultured on an air-liquid interface by discarding the apical medium and adding PneumaCult-ALI Basal Medium with supplements (Stemcell), 4 μg/ml heparin (Stemcell), 0.48 μg/ml hydrocortisone (Stemcell), and 100 μg/ml streptomycin and 100 U/ml penicillin (Lonza) to the basal compartment to induce differentiation. Cells were differentiated for 4 weeks on this air-liquid interface. Prior to experiments, the cells were cultured for 3 days in PneumaCult-ALI Basal Medium without heparin, hydrocortisone, and antibiotics (63).

### Viruses

SARS-CoV-2 isolate BetaCoV/Munich/BavPat1/2020 (European Virus Archive no. 026V-03883) was kindly provided by Prof. C. Drosten (Charité-Universitätsmedizin, Berlin Institute of Virology, Berlin, Germany) and was initially cultured in Vero E6 cells up to three passages in the laboratory of Prof. B. Haagmans (Viroscience Department, Erasmus Medical Center, Rotterdam, The Netherlands). To prepare virus stock, Vero FM cells were infected with passage 3 stock at an MOI of 0.01 in virus infection medium [DMEM (Gibco) containing 2% FCS (Sigma-Aldrich), 20 mM HEPES buffer (Gibco), 100 μg/ml streptomycin (Gibco), 100 IU/ml penicillin (Gibco). At 48 hpi cell culture supernatant was harvested, centrifuged at 4700 x *g* for 10 min at 4°C to remove cellular debris, filtered through a 0.2 μm syringe filter (Whatman) and stored as 100 μl aliquots at −80°C.

SARS-CoV-2 isolate CoV-2/human/Nijmegen/1/2020 was isolated from an oro-nasopharyngeal swab of a 65-year old male COVID-19 patient hospitalized at Radboud University Medical Center in May 2020, collected as part of an ongoing study on COVD-19 infectiousness and viral kinetics. The study was approved by the Committee on Research Involving Human Subjects Arnhem Nijmegen (CMO NL2020-6517) and conducted according to the principles of the Declaration of Helsinki (last updated 2013) and in accordance with the Medical Research Involving Human Subjects Act (Dutch: WMO). Verbal consent was provided at inclusion and separate verbal consent was obtained for the use of isolated virus in additional experiments.

The nasal swab was initially stored in virus transport medium [Hank’s Balanced Salt Solution (Gibco) containing 2% FCS (Sigma-Aldrich), 100 μg/ml gentamycin (Gibco) and 0.5 μg/ml amphotericin-B (Gibco)]. Vero FM cells seeded in 24-well plates were infected in duplicate with 100 μl 2-fold serial dilutions of patient material in a total volume of 200 μl using virus isolation medium [DMEM (Gibco) containing, 20 mM HEPES buffer (Gibco), 100 μg/ml streptomycin (Gibco), 100 IU/ml penicillin (Gibco), 1% amphotericin B (Gibco)]. After 1 h adsorption, viral inoculum was discarded, cells were washed with PBS, and 500 μl virus isolation medium containing 2% FCS was added. 100 μl of supernatant was collected at 2, 4 and 6 days post-infection, subjected to RNA isolation and RT-qPCR to detect SARS-CoV-2 RNA. The rest of the supernatants were stored at −80°C. Supernatant from which a positive signal was obtained in RT-qPCR was titrated by conventional plaque assay and was cultured in Vero FM cells at MOI 0.01 to obtain passage 1 working stocks. The near full-length viral sequence of the isolate was deduced by amplicon-based next-generation sequencing, supplemented with Sanger sequencing to fill few gaps in the obtained sequence. The sequence will be deposited in GenBank.

### Virus titration

Vero E6 cells were seeded onto 12-well plates at a density of 5 × 10^5^ cells/well. At 24 h post-seeding, cells were washed twice with PBS and infected with 200 μl 10-fold serial dilutions of the virus. After 1 h adsorption, the inoculum was discarded, cells were washed with PBS and overlay medium containing Minimum essential medium (MEM, Gibco), 2% FCS (Sigma-Aldrich), 20 mM HEPES buffer (Gibco), 0.75% carboxymethyl cellulose (Sigma-Aldrich), 100 μg/ml streptomycin and 100 IU/ml penicillin (Gibco) was added onto the cells. At 48 hpi, the medium was discarded, cells were washed with PBS, and stained with 0.25% crystal violet solution containing 4% formaldehyde for 30 minutes. Plates were then washed with double-distilled water, dried and plaques were counted.

### Infection of primary nasal epithelial cells

Differentiated primary nasal epithelial cells seeded onto 0.4 μm pore polyester membrane inserts (Corning) were infected with SARS-CoV-2 BavPat1 isolate at MOI 10 at both the apical and basolateral surfaces. At 2 hpi, the inoculum was discarded, cells were washed with PBS and fresh PneumaCult-ALI Basal Medium (Stemcell) without heparin, hydrocortisone and antibiotics was added to the basolateral compartment. At the desired time point, 200 μl of pre-warmed medium was added to the apical surface of the cells and incubated for 10 min at 37°C. Apical and basolateral supernatants were collected for titration or RNA isolation. Intracellular RNA was isolated using RNA-Solv reagent (Omega Bio-tek) according to manufacturer’s protocols, using glycogen during precipitation.

### Antiviral assay

BBR (Sigma-Aldrich) and OLX (Selleck Chemicals) were dissolved in DMSO at a stock concentration of 10 mM. Vero E6 cells were seeded onto 24-well plates at a density of 1.5 × 10^5^ cells/well. 24 h post-seeding, cells were washed twice with PBS and infected with the SARS-CoV-2 BavPat1 or Nijmegen1 isolate at MOI 0.01 in the presence of eight concentrations of BBR (150 μM – 1.2 μM) or OLX (5 μM – 0.04 μM) in a two-fold dilution series. As a negative control, SARS-CoV-2 infection in the presence of 0.1% DMSO, corresponding to the DMSO concentration in cells treated with 10 μM of compound. At 1 hpi, virus inoculum was discarded, cells were washed with PBS and replaced with infection medium containing the same concentrations of the inhibitors. At 24 hpi, cell culture supernatants were collected and stored at −80°C for plaque titration. RNA was isolated from cells or 100 μl cell culture supernatant using RNA-Solv reagent (Omega Bio-Tek) and precipitated in the presence of glycogen.

Differentiated primary nasal epithelial cells, seeded onto 0.4 μm pore polyester membrane inserts (Corning), were infected with SARS-CoV-2 BavPat1 isolate at MOI 10 at both the apical and basolateral surfaces in the presence of a two-fold dilution series of BBR (4.7 μM – 150 μM) or OLX (0.08 μM - 5 μM). At 2 hpi, the inoculum was discarded, cells were washed with PBS and fresh PneumaCult-ALI Basal Medium without heparin, hydrocortisone, and antibiotics, containing the same concentrations of BBR or OLX was added to the basolateral compartment. SARS-CoV-2 infection in the presence of 0.1% DMSO was used as a negative control. At 72 hpi, 200 μl of pre-warmed medium was added to the apical surface of the cells and incubated for 10 min at 37°C and 200 μl was collected for RNA isolation. The effective concentration of compound that reduced viral levels by 50% (EC_50_) was estimated by four parameter logistic regression, using Graphpad Prism (version 5.0). The selectivity index (SI) was defined as CC_50_/EC_50_.

### Cell viability assay

Vero E6 cells seeded in 96-well plates at a density of 3 × 10^4^ cells/well were treated with the same concentrations of BBR and OLX used in the antiviral assay in the absence of SARS-CoV-2 infection. Cells treated with 0.1% DMSO were used as a negative control. At 24 h post-treatment, cell viability was assessed using the Cell Titer Glo 2.0 kit (Promega) according to the manufacturer’s instructions. Luminescence was detected on the Victor Multilabel Plate Reader (Perkin Elmer). The 50% cytotoxicity concentration (CC_50_) value was estimated by four parameter logistic regression of the data, using Graphpad Prism (version 5.0).

### Time-of-addition assay

Vero E6 cells were seeded in 24-well plates at a density of 1.5 × 10^4^ cells/well. At 24 h post-seeding, cells were washed twice with PBS and infected with SARS-CoV-2 BavPat1 isolate at MOI 1. At indicated time points during the course of the experiment, 20 μM BBR, 0.25 μM OLX, or 0.1% DMSO was added. Cell culture supernatants were collected at 10 hpi and infectious viral titers were analyzed estimated by plaque assay.

### RT-qPCR

TaqMan Reverse Transcription reagent and random hexamers (Applied Biosystems) were used for cDNA synthesis. Semi-quantitative real-time PCR was performed using GoTaq qPCR Master Mix (Promega) using primers targeting the SARS-CoV-2 E protein gene (forward primer, 5′-ACAGGTACGTTAATAGTTAATAGCGT-3′; reverse primer, 5′-ACAGGTACGTTAATAGTTAATAGCGT-3’). A standard curve of a plasmid containing the E gene qPCR amplicon was used to convert Ct values to relative genome copy numbers. Human and African green monkey (*Chlorocebus sabaeus*) β-actin were used as housekeeping genes for normalization (Human β-actin, forward primer 5′-CCTTCCTGGGCATGGAGTCCTG-3′ and reverse primer 5′-GGAGCAATGATCTTGATCTTC-3′; African green monkey β-actin, forward primer 5′-ATTGGCAATGAGCGGTTCC-3′ and reverse primer 5′-CTGTCAGCAATGCCAGGGTA-3′).

### Immunofluorescence staining

Primary nasal epithelial cells were mock-infected with PneumaCult-ALI Basal Medium for 2 h or infected with SARS-CoV-2 BavPat1 isolate at MOI 10. Vero E6 cells were infected with SARS-CoV-2 BavPat1 isolate at MOI 0.01 or mock-infected for 1 h at 37°C, after which cells were washed with PBS. Primary nasal epithelial cells were fixed with 4% formaldehyde at 0, 24, 48, 72, and 96 hpi. Vero E6 cells (Fig. 4) were fixed at 24 hpi. Cells were permeabilised with 0.1% Triton X-100 for 5 minutes, blocked with 2% Normal Serum Block (BioLegend), 1% BSA, and 0.0005% Triton X-100 in PBS for 30 minutes. Primary nasal epithelial cells and Vero E6 cells (Fig. 4 and Fig. S3) were stained for 1 h with 0.01 mg/ml rabbit anti-SARS-CoV-2 Spike S1 subunit (clone#007, Sino Biologicals) and 0.03 mg/ml mouse anti-Tubulin IV (cloneONS.1A6, Sigma) or mouse anti-Muc5AC (clone 45M1, Invitrogen). Subsequently, the cells were stained for 1 h with 0.01 mg/ml goat anti-mouse Dylight 488 (Biolegend), 0.01 mg/ml donkey anti-rabbit AF555 (Biolegend), and a 1:2000 dilution of phalloidin-iFluor 647 reagent (Abcam). Vero E6 cells (Fig. S1) were stained for 1 h with anti-SARS-CoV Spike S1 subunit human IgG1 (BEI Resources) and mouse anti-dsRNA J2 monoclonal antibody (Scicons). Goat anti-mouse Alexa Fluor 568 (Invitrogen) and goat anti-human Alexa Fluor 488 (Invitrogen) were used at 1:500 dilutions for secondary staining. The polyester membrane containing the nasal epithelial cells was cut out of the insert with a scalpel and placed on a microscopy slide, embedded in ProLong Diamond Antifade Mountant with DAPI (Invitrogen), and covered with a coverslip. Vero E6 that had been grown on coverslips were embedded in the same way. Fluorescent images were made using a Leica Dmi8 microscope (Leica) with 20× and 100× objectives and Leica CFC7000 GT camera using LAS X 3.4.2 software. Confocal images for Vero E6 cells in Fig. S1B were captured using a Zeiss LSM9000 microscope with a 63× oil objective and processed using FIJI software.

### Cytokine and chemokine analysis

Primary nasal epithelial cells cultured on an air-liquid interface were infected with SARS-CoV-2 (BavPat1 isolate) at MOI 10 or mock infected. Samples were taken from the basolateral medium over time and the concentration of chemokines and cytokines was determined using the LEGENDplex Human Anti-Virus Response Panel (13-plex) and the LEGENDplex Human Proinflammatory Chemokine Panel (13-plex) on a FACSCanto II flow cytometer (BD). Samples were run in duplicate and measured at two different dilutions.

## Acknowledgments

We thank Jordy Coolen (Radboud University Medical Center) for help with sequencing the SARS-CoV-2 Nijmegen1 strain. We thank Christian Drosten (Charité-Universitätsmedizin Berlin, Institute of Virology, Berlin, Germany) and Bart Haagmans (Viroscience Department, Erasmus Medical Center, Rotterdam, The Netherlands) for providing SARS-CoV-2 (isolate BetaCoV/Munich/BavPat1/2020), and Kjerstin Lanke (Radboud University Medical Center) for providing the plasmid used as standard curve in the qPCR. The following reagent was obtained through BEI Resources, NIAID, NIH: Monoclonal Anti-SARS Coronavirus Recombinant Human IgG1, Clone CR3022 (catalog number NR-52392). This work was financially supported by a Research Grant from the Human Frontiers Science Foundation (grant number RGP0007/2017) and a VICI grant from the Netherlands Organization for Scientific Research (grant number 016.VICI.170.090) to R.P. van Rij.

