## Supplemental figures for "Berberine and obatoclax inhibit SARS-CoV-2 replication in primary human nasal epithelial cells in vitro"

### Supplementary Figures

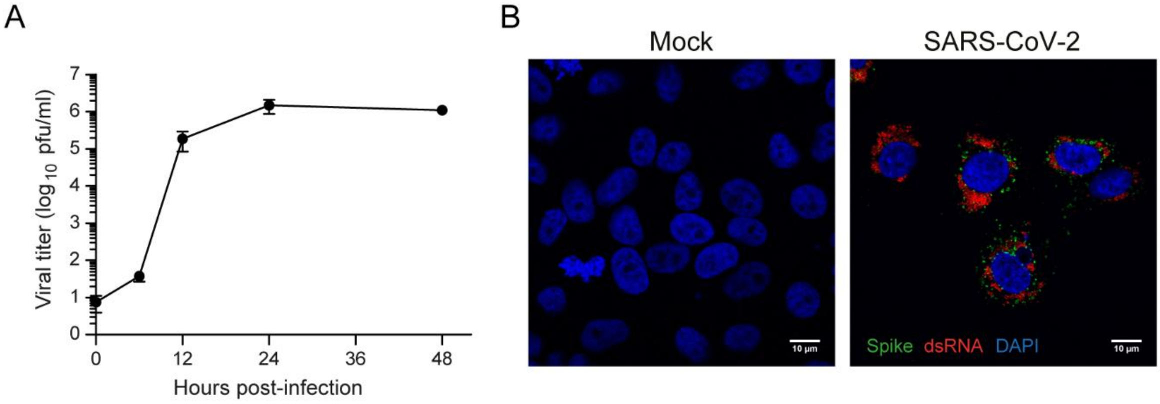


**Fig. S1. SARS-CoV-2 replication kinetics in Vero E6 cells. (A)** Viral infectious titers from Vero E6 cell infected with SARS-CoV-2 (BavPat1 isolate, MOI=0.01) in culture supernatants over time. Error bars represent SD of n = 2 replicates. **(B)** Indirect immunofluorescence staining of mock-infected or SARS-CoV-2 infected Vero E6 cells (MOI=0.01). The presence of dsRNA (red) and SARS-CoV-2 Spike protein S1 subunit (green) was assessed at 24 hpi. Nuclei were stained with DAPI (blue). Bar represents 10 µm.

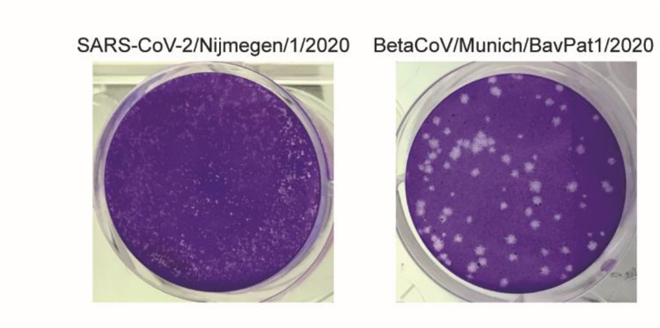


**Fig. S2. Plaque phenotype of SARS-CoV2 Nijmegen1 isolate.** Plaque phenotype in Vero E6 cells of the SARS-CoV-2 Nijmegen1 (left) and BavPat1 (right) isolates used in this study. Differences in plaque phenotypes may be due to the different passage history of both isolates in the Vero cell line.

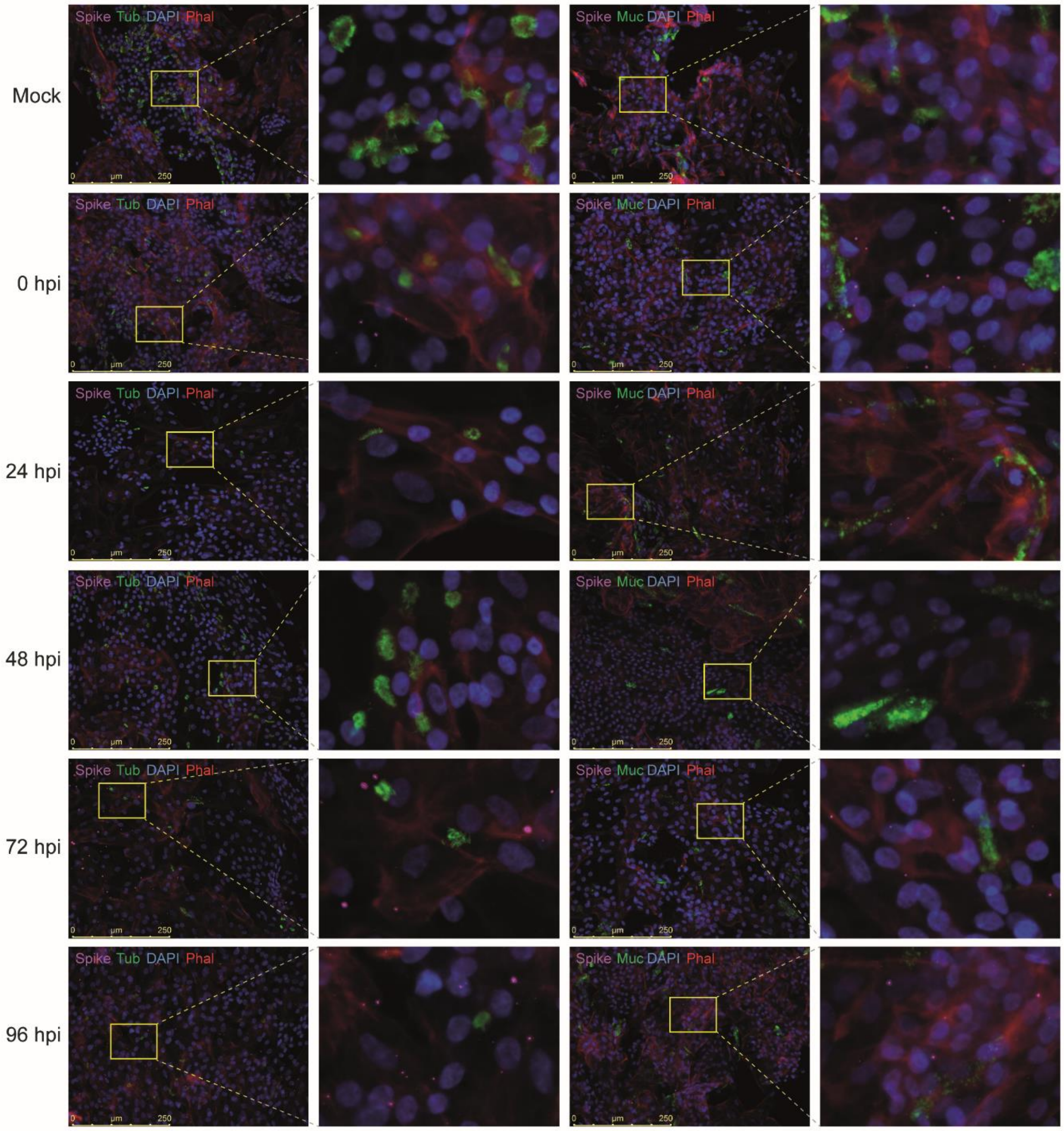


**Fig. S3. SARS-CoV-2 spike protein expression in primary nasal epithelial cells.** Indirect immunofluorescence staining of mock-infected or SARS-CoV-2 infected primary nasal epithelial cells (MOI=10), fixed at the indicated time points post-infection. The presence of SARS-CoV-2 Spike protein subunit S1 (pink) was assessed in cells stained with anti-tubulin IV (ciliated cells, green, left panels) or anti-Muc5AC (goblet cells, green, right panels), DAPI (nucleus, blue), and phalloidin (F-actin, red). Bars represent 250 µm. Yellow boxes indicate the regions shown at higher magnification in the right panels.

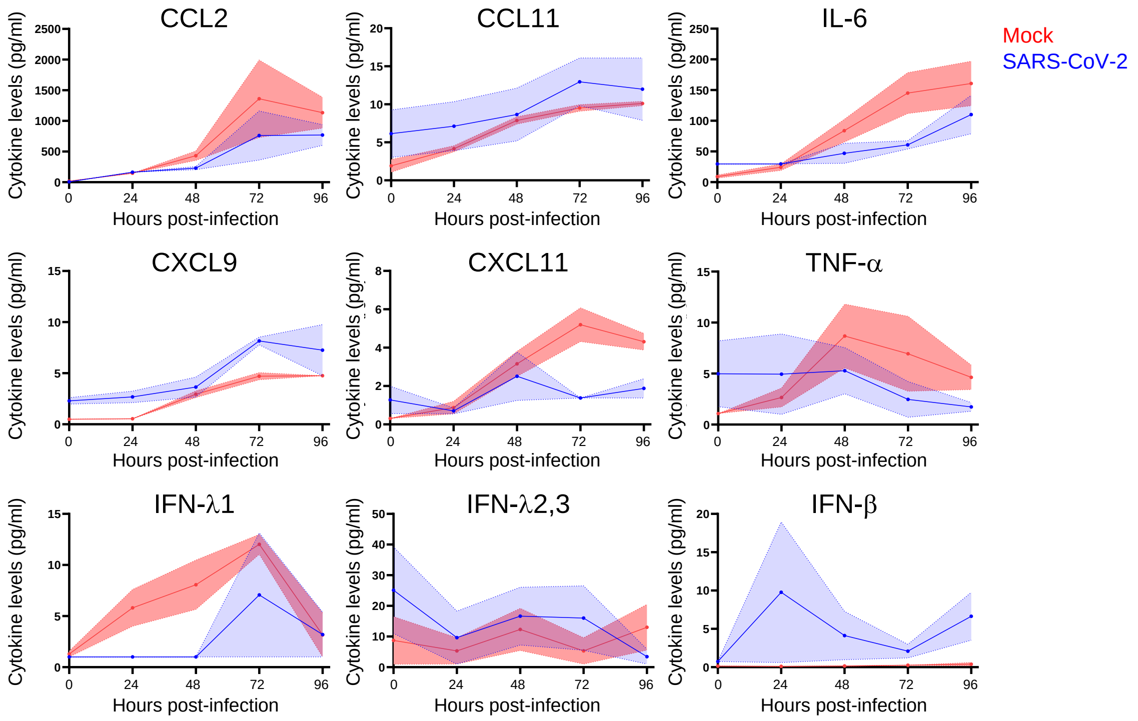


**Fig. S4. Cytokine and chemokine production from SARS-CoV-2 infected primary nasal epithelial cells.** Primary nasal epithelial cells, cultured on an air-liquid interface, were infected with SARS-CoV-2 (BavPat1 isolate) at an MOI of 10 or mock infected. At the indicated time points, medium from the basolateral compartment was harvested and the production of the indicated cytokines and chemokines was analyzed by a bead-based immunoassay. Means and SEM (shading) of n = 2 replicates are shown.
